# Cell-free identification of *de novo* designed minibinders as building blocks for targeted protein degraders

**DOI:** 10.64898/2026.11.22.768393

**Authors:** Chandra K. Bandi, Erik J. Bidstrup, Darianna Colina, Mehman I. Bunyatov, Thomas C. Donahue, Giancarlo Chuquitarco, William K.M. Lai, Matthew P. DeLisa

**Author notes:** Address correspondence to: Matthew P. DeLisa, Robert Frederick Smith School of Chemical and Biomolecular Engineering, Cornell University, Ithaca, NY 14853., Wiliiam KM Lai, Department of Biochemistry, Jacobs School of Medicine and Biomedical Sciences, State University of New York at Buffalo, Buffalo, NY 14203. These authors contributed equally to this work (Chandra K. Bandi, Erik J. Bidstrup, Darianna Colina).

## Abstract

Targeted protein degradation is a powerful strategy for manipulating protein abundance and function, but developing protein-based degraders typically requires laborious screening and optimization of target-specific binders. Recent advances in *de novo* protein design offer a promising route to accelerate degrader development, although computationally designed binders often exhibit variable binding activity and require extensive experimental validation. Here, we developed a rapid screening platform that couples cell-free protein synthesis (CFPS) with enzyme-linked immunosorbent assays (ELISA) to identify functional miniprotein binders directly from computational designs. Using β-catenin as a model target, we screened 93 *de novo* designed minibinders for expression and binding activity in less than three days, identifying six candidates with half-maximal effective concentrations below 500 nM. Two lead minibinders exhibited binding affinities of 299 and 335 nM and, without further affinity maturation, were fused to an engineered E3 ubiquitin ligase to generate functional protein degraders. These constructs reduced cytosolic β-catenin levels by approximately 80% and 60%, respectively, and suppressed β-catenin-dependent transcriptional activity in mammalian cells. These findings demonstrate that moderate-affinity, computationally designed minibinders can support effective targeted protein degradation without iterative binder optimization. More broadly, our integrated design and screening workflow provides a rapid and accessible approach for converting *de novo* protein binders into functional intracellular degraders.

## INTRODUCTION

Targeted protein degradation has emerged as a powerful strategy for perturbing cellular processes, investigating protein function, and developing new therapeutic modalities ^1^. However, the vast diversity of the proteome presents a significant challenge for developing degraders that selectively target individual proteins, particularly those considered undruggable because they lack suitable small-molecule binding pockets or because ligand binding does not necessarily disrupt their biological activity ^2,3^. Unlike conventional small-molecule inhibitors, which typically require sustained target occupancy to suppress protein function, targeted degradation offers a means to eliminate proteins without directly inhibiting their activity ^4^. One strategy involves engineering E3 ubiquitin ligases with target-specific protein or peptide binders that recruit proteins of interest for ubiquitination and subsequent proteasomal degradation ^5–7^.

Recent advances in *de novo* protein design have enabled the generation of target-specific binders that could greatly expand the repertoire of proteins amenable to targeted degradation ^8–10^. Among these, miniproteins are particularly attractive because of their small size, structural stability, and ability to recognize defined protein epitopes. Computationally designed miniproteins have been used as standalone binders to modulate cellular processes and enable *in situ* imaging ^11–17^ or fused to other proteins to redirect endosomal trafficking ^18^ and mediate cytotoxic targeting of CD8+ CAR-T cells ^19^. However, existing *de novo* binder design workflows typically rely on yeast surface display, biolayer interferometry (BLI), and iterative optimization to identify high-affinity binders, making experimental validation costly and time-consuming ^20^. Alternatively, diffusion-based peptide design methods have enabled the generation of moderate-affinity “guide” peptide binders without requiring target structural information ^21^. Approaches including SaLT&PepPr, PepPrCLIP, and PepMLM have yielded peptide binders that, when fused to E3 ligases, effectively degraded diverse intracellular protein targets, including the oncogenic signaling protein β-catenin, despite possessing only mid-nanomolar binding affinities ^22–25^. However, these workflows relied on screening peptides after fusion to E3 ligases, limiting screening throughput and complicating independent characterization of binder activity. Moreover, unlike structure-guided miniprotein design, these approaches do not provide precise control over the targeted binding epitope.

Cell-free protein synthesis (CFPS) offers a promising approach to overcome these screening limitations by enabling rapid, parallel expression and functional characterization of computationally designed binders. CFPS can produce proteins directly from linear DNA expression templates within hours, making it well suited for high-throughput screening ^26–28^. This capability has enabled the simultaneous evaluation of hundreds of protein variants, including iterative enzyme optimization through deep learning-guided design-build-test cycles ^27^, and characterization of previously identified binders using *in situ* BLI ^29–31^, AlphaLISA ^26^, and two-hybrid assays ^32^. However, these approaches have not been widely applied to the initial screening of *de novo* designed miniproteins, and some require high-affinity interactions for reliable detection. Integrating computational binder design with CFPS-based screening could therefore enable the rapid identification of moderate-affinity miniproteins suitable for targeted degradation without extensive binder optimization.

Here, we developed an integrated workflow combining *de novo* miniprotein design, CFPS, and enzyme-linked immunosorbent assays (ELISA) to rapidly identify functional binders for targeted protein degradation. Using the AlphaFold2-predicted structure of the historically difficult-to-drug signaling protein β-catenin ^33^, we generated miniprotein binders (minibinders) against a defined epitope using RFdiffusion ^34^ and screened 93 designs for expression and binding activity directly from CFPS reactions programmed with linear DNA templates. This approach identified six binders with half-maximal effective concentrations below 500 nM, including two lead candidates with binding affinities of 299 and 335 nM. Without further affinity maturation, these binders were fused to an engineered E3 ubiquitin ligase and shown to reduce cytosolic β-catenin levels by approximately 80% and 60%, respectively, while suppressing β-catenin-dependent transcriptional activity. Together, these results establish a rapid and accessible workflow for converting computationally designed miniproteins into functional protein degraders and demonstrate that moderate-affinity binders can support effective intracellular protein degradation.

## RESULTS

### De novo design of minibinders against a defined epitope on β-catenin

We sought to determine whether minibinders generated from a single round of *de novo* design, without subsequent affinity maturation, could serve as targeting domains for functional protein degraders. To test this, we established a four-step workflow comprising computational backbone and sequence design, CFPS from linear DNA templates, ELISA-based screening for expression and target binding, and fusion of selected binders to an E3 ubiquitin ligase for functional evaluation in mammalian cells (**Fig. 1a**). We selected human β-catenin (CTNNB1) as a model target because of its established role key driver of aberrant Wnt signaling in cancer ^35^, the challenges associated with selectively inhibiting its biological activity using conventional small molecules ^33^, and previous demonstrations of peptide-directed degradation that provide a benchmark for the binding affinities required for effective protein knockdown ^22–24^.

**Figure 1.**
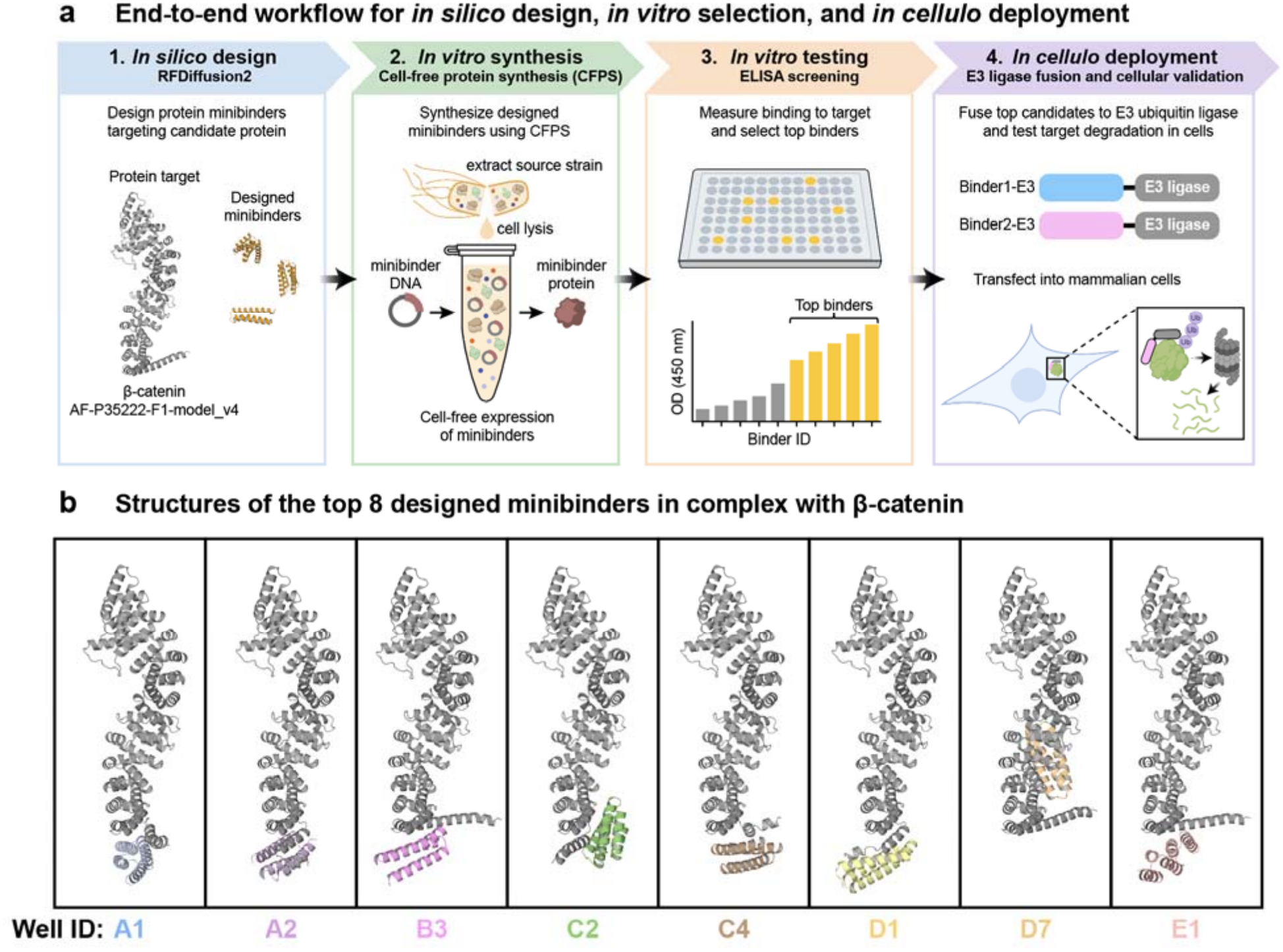
*De novo* design and screening of minibinders targeting β-catenin. (a) Schematic of the four-step workflow comprising computational binder design, cell-free protein synthesis (CFPS), ELISA-based screening, and cellular validation. Miniprotein binders (minibinders) targeting human β-catenin (CTNNB1) were designed using RFdiffusion and an AlphaFold2-predicted target structure. Designed binders were synthesized from DNA templates using an *E. coli*-based CFPS system and screened for target binding by ELISA. Selected binders were fused to an E3 ubiquitin ligase and expressed in mammalian cells to evaluate targeted β-catenin degradation. (b) Predicted structures of the eight selected minibinders (colored) in complex with β-catenin (gray), labeled by synthesis-plate well ID.

To evaluate whether computationally predicted target structures could support this single-round design strategy, we used the AlphaFold2 model of full-length human β-catenin (AF-P35222-F1, v4) as the starting template. Because the intrinsically disordered N- and C-terminal regions were predicted with low confidence, we removed these segments based on sliding-window smoothing of per-residue pLDDT scores, retaining the structured armadillo repeat core (see Materials and Methods for details). Using RFdiffusion ^34^, we generated 10,000 candidate binder backbones comprising 70–100 residues. ProteinMPNN was then used to design sequences for each backbone, followed by prediction of the corresponding binder–target complexes to calculate pLDDT, interface predicted template modeling (ipTM) scores, and interchain predicted aligned error (PAE). The resulting designs consisted of compact α-helical bundles characteristic of computationally designed miniproteins. We selected the 96 top-ranked candidates for DNA synthesis, of which 93 met the vendor’s synthesis requirements. Predicted structures of the nine minibinders subsequently selected for purification are shown in **Fig. 1b**.

### CFPS-enabled screening of miniprotein expression and binding activity

To rapidly evaluate the expression and β-catenin-binding activity of *de novo* designed minibinders, we obtained synthetic DNA fragments encoding the 93 top-ranked designs. Each fragment was PCR-amplified to generate a linear expression template (LET) for CFPS ^36^. Following 16 h of CFPS, reactions were diluted and directly analyzed by ELISA to assess relative minibinder expression (**Fig. 2a**) and binding to immobilized β-catenin (**Fig. 2b**). Although many minibinders exhibited robust expression, only about 10% showed appreciable β-catenin binding in our screen (**Supplementary Fig. 1**), which is in line with the expected hitrate for RFDiffusion-guided minibinder design on a complex protein target ^34^.

**Figure 2.**
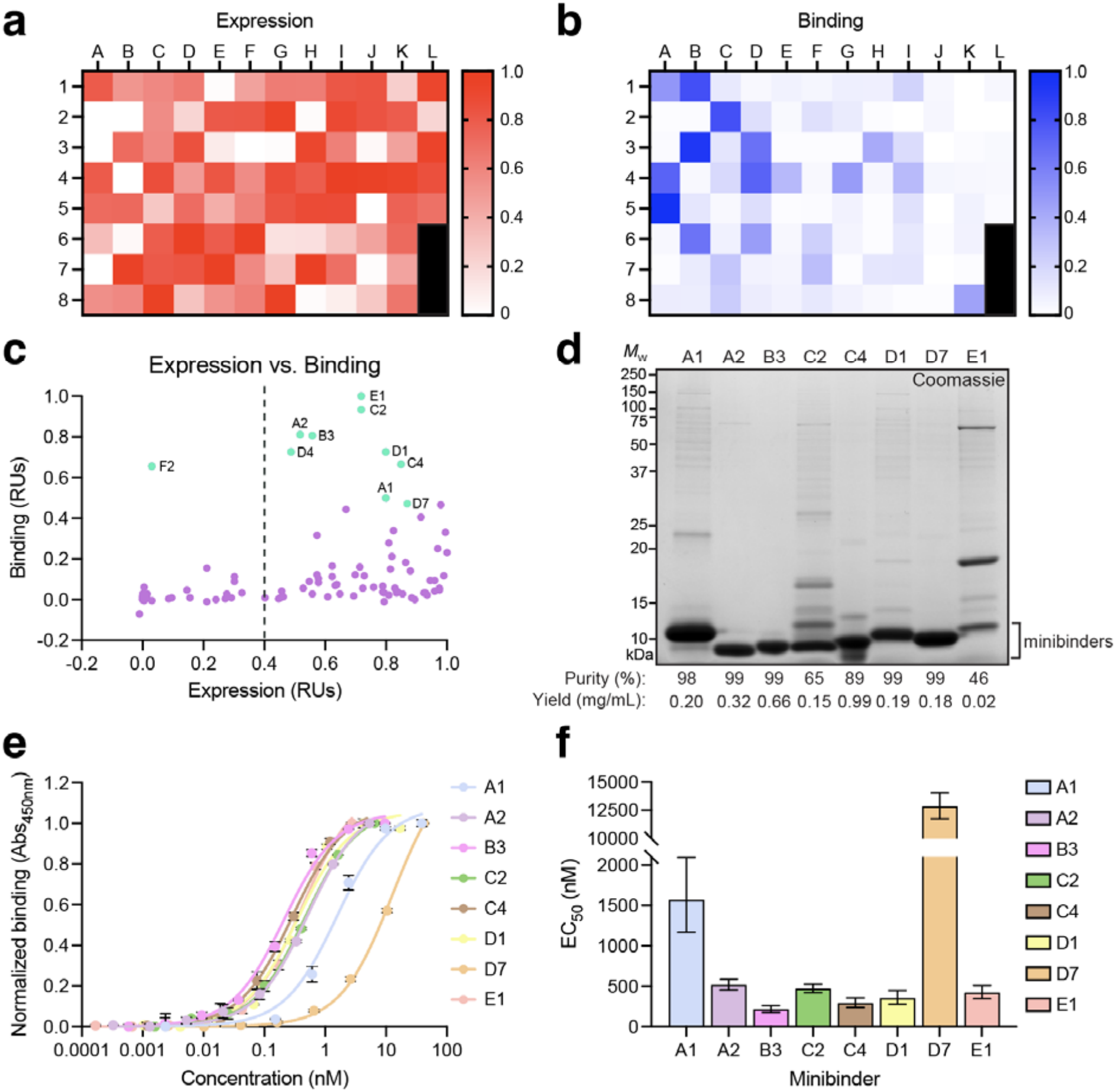
CFPS-enabled screening and validation of *de novo* designed β-catenin minibinders. (a) Heatmap of relative minibinder expression measured by direct ELISA using diluted CFPS reactions immobilized on high-binding plates and detected with an anti-His-HRP antibody. (b) Heatmap of relative minibinder binding measured by ELISA using diluted CFPS reactions and immobilized recombinant β-catenin, with bound minibinders detected using ExtrAvidin-HRP. For both assays, background-subtracted absorbance at 450 nm was normalized to the maximum signal, and values represent the mean of three independent CFPS reactions. Black area indicates blank wells. (c) Scatter plot comparing relative expression and binding signals for all 93 minibinder designs. The 10 highest-scoring binders are highlighted in teal and labeled by synthesis-plate well ID. The dashed line indicates the minimum expression threshold of 0.4 relative units (RUs) used for candidate selection. (d) Coomassie-stained SDS-PAGE gel of eight minibinders purified following recombinant expression in *E. coli*, with corresponding protein purity (%) and yield (mg/mL). Purity was determined by densitometry, and yield was calculated from total protein concentration measured by BCA assay and the corresponding purity. (e) ELISA binding curves for purified, biotinylated minibinders immobilized on high-binding plates and probed with recombinant β-catenin. Data represent the mean ± SD of two technical replicates. Individual binding curves are provided in **Supplementary Fig. 3**. (f) EC50 values with 95% confidence intervals determined from the binding curves in (e).

Using this knowledge, we next selected candidates for further characterization based primarily on their binding activity in the CFPS-ELISA screen. Out of the 93 total designs, the 10 highest-scoring minibinders were initially selected based on relative binding signals, after which a minimum expression threshold of 0.4 relative units (RUs) was applied to identify nine candidates with sufficient expression and strong binding activity (**Fig. 2c**). Notably, F2 exhibited strong β-catenin binding in our screen but fell below the expression threshold and was excluded from further characterization. To independently validate these candidates, we cloned their coding sequences into pET28a expression vectors, expressed them in *E. coli*, and purified the resulting proteins using StrepXT affinity chromatography. Protein purity was assessed by SDS-PAGE and Coomassie staining, while yields were determined by bicinchoninic acid (BCA) assay. Eight of the nine minibinders were successfully purified in quantities sufficient for further characterization (**Fig. 2d**), whereas D4 exhibited poor recovery following affinity purification (**Supplementary Fig. 2**). The purified minibinders were biotinylated, immobilized on ELISA plates, and assayed for binding to recombinant β-catenin to determine binding curves (**Fig. 2e** and **Supplementary Fig. 3**) and half-maximal effective concentrations (EC_50_) (**Fig. 2f**). A total of six of the eight purified minibinders exhibited EC_50_ values below 500 nM, corresponding to a 75% hit rate among experimentally characterized candidates and 67% of the nine initially selected designs. Notably, EC_50_ values showed a moderate correlation with relative binding signals from the initial CFPS-ELISA screen (**Supplementary Fig. 4**), indicating that the screening platform could identify moderate-affinity binders despite potential differences in expression levels among designs.

**Figure 3.**
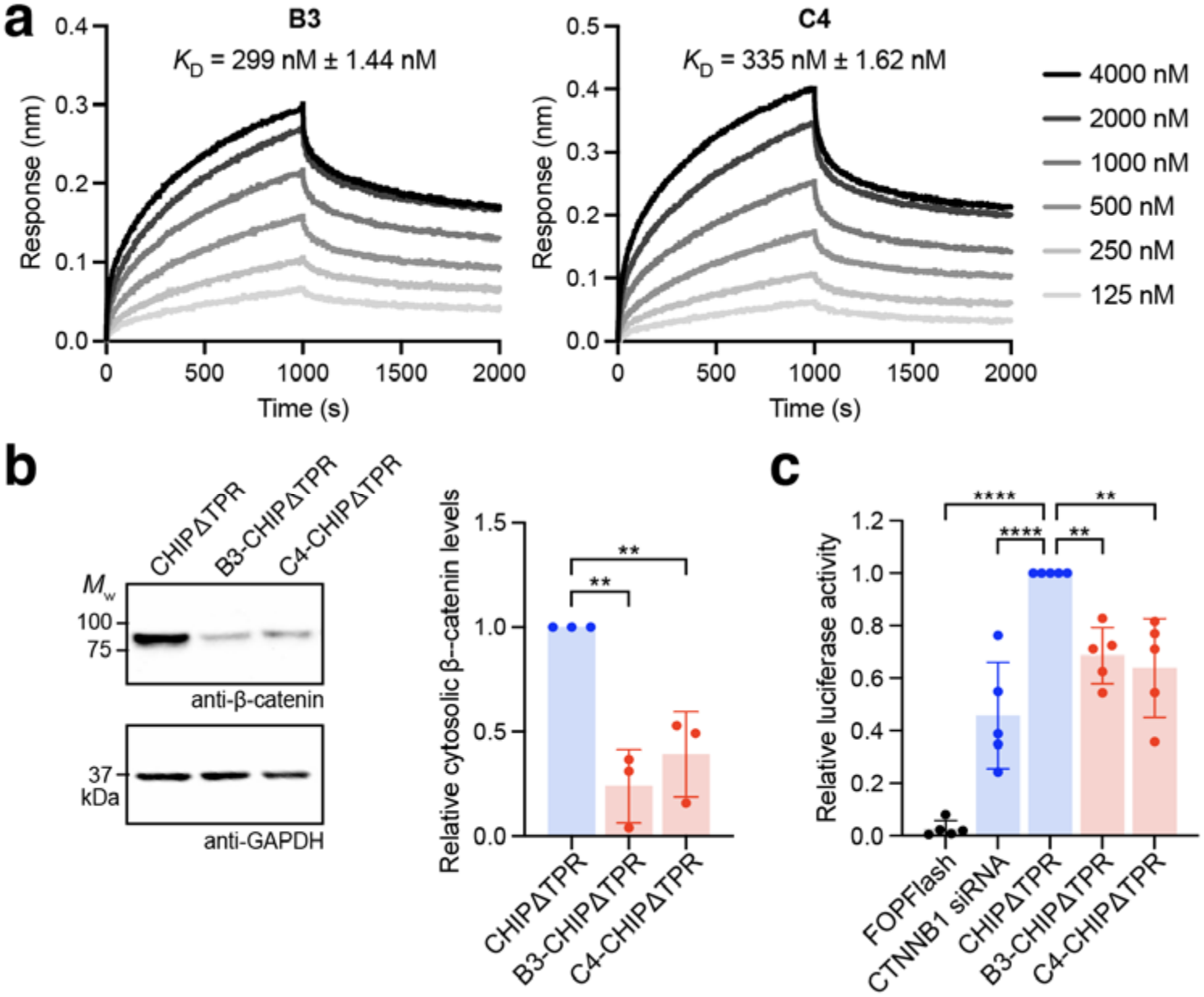
Top identified minibinder degraders exhibit robust binding and enable functional β-catenin degradation. (a) BLI sensorgrams for binding of immobilized minibinders B3 (top) and C4 (bottom) to recombinantly produced β-catenin. Data were fitted to a 1:1 monovalent analyte model to obtain *K*_D_ ± 95% confidence interval using GraphPad Prism for macOS (version 11.1.0). (b) Representative immunoblots and quantification of cytosolic β-catenin levels in HEK293T cells 48 h after transfection with plasmids encoding CHIPΔTPR, B3–CHIPΔTPR, or C4–CHIPΔTPR. Densitometri analysis was performed using ImageJ software (version 1.37v). GAPDH was used as a loading control, and cytosolic β-catenin levels were normalized to the CHIPΔTPR control. (c) Relative TOPFlash luciferase activity in HEK293T cells 48 h after transfection with plasmids encoding CHIPΔTPR, B3-CHIPΔTPR, C4-CHIPΔTPR, or CTNNB1 siRNA with FOPFlash included as a negative control. Luciferase activity was normalized to the CHIPΔTPR control. Data are presented as mean of biological replicates (*n* = 3 for panel b; *n* = 5 for panel c) ± SD. Statistical significance was assessed using one-way ANOVA. Calculated *p*-values are represented as follows: *, *p* < 0.05; **, *p* < 0.01; ****, *p* < 0.0001; ns, not significant.

### Construction and characterization of minibinder-guided degraders targeting β**-catenin**

We next selected minibinders B3 and C4, which exhibited the strongest β-catenin binding based on ELISA-derived EC_50_ values, for further characterization and development as targeted protein degraders. We measured their binding affinities by biolayer interferometry (BLI) using an Octet RH16 instrument (**Fig. 3a**), yielding equilibrium dissociation constants (*K*_D_) of 299 nM for B3 and 335 nM for C4, consistent with their ELISA-derived EC_50_ values (**Fig. 2e**). To convert these minibinders into targeted protein degraders, we genetically fused each to a truncated form of the human E3 ubiquitin ligase C-terminus of Hsp70-interacting protein (CHIP) lacking its native tetratricopeptide repeat (TPR) substrate-binding domain (CHIPΔTPR) ^37^. The resulting fusion constructs, designated minibinder-guided degraders B3 and C4, were cloned into the mammalian expression vector pcDNA3.

To evaluate the degradation activity of the minibinder-guided degraders, we transiently transfected HEK293T cells with pcDNA3 constructs encoding B3 or C4 fused to CHIPΔTPR, or CHIPΔTPR alone as a control lacking the target-binding domain. After 48 h, cytosolic β-catenin levels were assessed by immunoblotting. We focused on the cytosolic pool because its accumulation promotes nuclear translocation of β-catenin, where it interacts with TCF/LEF transcription factors to activate genes involved in oncogenic signaling ^38^. Densitometry analysis revealed that degraders B3 and C4 significantly reduced cytosolic β-catenin levels by approximately 80% and 60%, respectively, relative to the CHIPΔTPR control (**Fig. 3b**).

To determine whether β-catenin degradation also reduced downstream transcriptional activity, we performed a TOPFlash reporter assay, which measures β-catenin-dependent transcription through tandem TCF/LEF-binding motifs upstream of a luciferase reporter gene ^39,40^. We compared the two minibinder-guided degraders to an siRNA control specific to the β-catenin gene, CTNNB1, which was reported previously ^24^. Consistent with the immunoblotting results, both degraders significantly reduced β-catenin transcriptional activity by approximately 30% and 35%, respectively, relative to the CHIPΔTPR control (**Fig. 3c**). Interestingly, no significant difference in TOPFlash reporter activity was observed between either minibinder-fused degrader and CTNNB1 siRNA, suggesting comparable effects on β-catenin dependent transcriptional activity. Together, these results demonstrate that minibinder-fused degraders B3 and C4 effectively reduce cytosolic β-catenin levels and suppress its downstream transcriptional activity.

## DISCUSSION

Here, we established a rapid and accessible CFPS-enabled workflow for identifying *de novo* designed minibinders and implementing them as guide domains for targeted protein degradation. The workflow enabled expression and binding screening of 93 computationally designed minibinders in less than three days, followed by independent validation of selected candidates. Of the nine minibinders selected for further characterization, eight were successfully purified, and six exhibited ELISA-derived EC_50_ values below 500 nM. Without additional affinity maturation, two of these minibinders were converted into functional protein degraders by fusion to the E3 ubiquitin ligase CHIPΔTPR. The resulting constructs reduced cytosolic β-catenin abundance by up to 80% and suppressed β-catenin-dependent transcriptional activity to levels not significantly different from those achieved with CTNNB1-targeting siRNA.

Despite substantial advances in computational protein design, the discovery of high-affinity *de novo* minibinders often requires screening large libraries of candidates followed by iterative optimization or affinity maturation ^41^. These approaches have enabled the development of potent binders against diverse protein targets and offer important advantages over existing methods for generative peptide binder design, particularly the ability to direct binding toward defined epitopes on a target protein ^34^. However, the high affinities typically sought in these efforts may not be necessary for applications involving targeted protein degradation. We hypothesized that moderate-affinity minibinders identified through smaller-scale screening could serve as effective targeting domains when coupled to an E3 ubiquitin ligase. This premise is consistent with the moderate binding affinities observed for native E3 ligase interactions (e.g., CHIP binds its native substrates Hsp70, Hsp90, and Hsc70 with *K*_d_ values of 0.3–2.3 μM) ^42^ and the catalytic, nonstoichiometric nature of ubiquitin-mediated protein degradation ^43^. Indeed, minibinders B3 and C4 bound β-catenin with equilibrium dissociation constants of 299 and 335 nM, respectively, comparable to the 250–430 nM affinities reported for peptide binders previously used to direct CHIP-mediated β-catenin degradation ^22,24^. These findings support the idea that high-affinity binding is not an absolute requirement for effective targeted protein degradation. Instead, degrader activity may depend on additional factors, including binding kinetics, target engagement, epitope accessibility, and the geometry of the resulting complex relative to sites available for ubiquitination ^44^.

The feasibility of using moderate-affinity binders motivated our development of a simplified CFPS screening strategy for evaluating a relatively small set of computational designs. Previous studies have established the utility of CFPS for rapid, purification-free screening of antibody fragments and nanobodies using AlphaLISA or biolayer interferometry ^26,29,30^. More recently, Capin et al. extended cell-free screening to *de novo* designed protein binders using a transcriptional two-hybrid assay, although interactions with dissociation constants above approximately 100 nM were not reliably detected ^32^. Unlike recent high-throughput workflows for screening computationally designed protein libraries that rely on plasmid cloning, bacterial transformation, cell-based expression, and subsequent protein purification ^45^, our approach combined LET-based CFPS with direct ELISA analysis to independently assess minibinder expression and binding without these additional steps. This enabled screening within 24 h of obtaining the expression templates, compared with the approximately one week reported for a recent high-throughput characterization campaign involving 96 computationally designed proteins ^45^.

Importantly, our workflow identified moderate-affinity minibinders that could be implemented as functional protein degraders without iterative affinity maturation. Of the nine candidates selected for further characterization, six subsequently exhibited EC_50_ values below 500 nM, demonstrating that the initial screen could identify binders with affinities suitable for further development as targeted protein degraders. Because LET generation, CFPS, and ELISA are readily scalable and compatible with automation, this strategy could support screening of substantially larger design libraries targeting multiple proteins or distinct epitopes within the same protein. The ability to direct *de novo* minibinder design toward specific structural epitopes could be particularly valuable for identifying binding sites that favor productive ubiquitination and efficient target degradation. Together, these features establish a streamlined, “single-shot” approach to minibinder discovery that enables rapid progression from *in silico* design to experimentally validated candidate binders in less than three days, without iterative affinity maturation. This strategy could accelerate the development of functional protein degraders and enable high-throughput investigations of protein function through targeted degradation.

We selected β-catenin as a model target because its distinct subcellular functions present an important challenge for therapeutic intervention. Whereas cytosolic accumulation and nuclear signaling contribute to oncogenic transcriptional programs, membrane-associated β-catenin plays an essential role in cadherin-mediated cell adhesion ^46,47^. Our minibinder-based degraders substantially reduced cytosolic β-catenin abundance and downstream transcriptional activity, although whether they selectively spare membrane-associated β-catenin remains to be determined. An additional limitation is that our experiments were performed in HEK293T cells rather than a cancer model dependent on aberrant β-catenin signaling. Future studies in β-catenin-dependent cancer cells, such as DLD1 ^24^, will be important for determining whether the observed reductions in β-catenin abundance and transcriptional activity translate into impaired tumor cell growth.

Together, our findings establish a rapid and accessible platform for converting *de novo* designed minibinders into functional protein degraders without iterative affinity maturation. By demonstrating that moderate-affinity binders can effectively direct targeted protein degradation, this work expands the utility of computational protein design beyond applications requiring exceptionally high binding affinities. The simplicity and scalability of this approach could enable the systematic development of programmable protein degraders against diverse targets, providing new tools for investigating protein function and advancing therapeutic discovery.

## MATERIALS AND METHODS

### *In silico* protein design

*De novo* protein binders targeting β-catenin were generated using RFdiffusion ^34^. The AlphaFold2-predicted β-catenin structure (AF-P35222-F1, version 4) ^48^ was trimmed to remove predicted intrinsically disordered regions at the N- and C-termini, with truncation boundaries determined using sliding-window smoothing of per-residue pLDDT scores. Design calculations specified target residues A100–200 and binder lengths of 70–100 amino acids (contigmap.contigs = [A100-200/0 70-100]), with residues A142, A143, A144, A145, A148, and A149 designated as binding hotspots. Residue numbering is relative to the processed input structure. A total of 10,000 designs were generated, and candidates were ranked by binder–target interchain predicted aligned error (PAE) ^41^, with the lowest-scoring designs selected for experimental characterization.

### Strains

*E. coli* strain DH5α was used for all plasmid construction in this study. *E. coli* strain BL21(DE3) (Novagen) was used for expression and purification of recombinant β-catenin. *E. coli* strain SHuffle T7 Express (NEB) was used to produce lysates for CFPS experiments.

### Plasmids and LETs

All DNA encoding the candidate minibinders were ordered as gBlocks (Twist Bioscience). These gBlocks were amplified using primers encoding the promoter and ribosome binding site (RBS) for LET production and used as the template for CFPS. The down-selected minibinder gBlocks were subsequently cloned into the pET28a(+) backbone using custom designed primers and Gibson assembly. Plasmid M57 pET28a-TEV-full-length human beta catenin (Addgene #17198) was used for expression of recombinant β-catenin in *E. coli*. For construction of minibinder-guided degrader plasmids, we used a pcDNA3-based plasmid containing an Esp3I restriction site immediately upstream of DNA encoding a flexible GSGSG linker followed by the CHIPΔTPR gene ^22,24^. The B3 and C4 minibinder gBlocks were cloned into the Esp3I-digested pcDNA3-CHIPΔTPR backbone using custom designed primers and Gibson assembly. Assembled constructs were used to transform DH5α cells and plated on Luria-Bertani (LB) agar supplemented with the appropriate antibiotic. All plasmids were isolated using a QIAprep Spin Miniprep Kit (Qiagen) and confirmed by DNA sequencing at the Genomics Facility of the Cornell Biotechnology Resource Center.

### CFPS-enabled minibinder screening

DNA gBlocks encoding candidate minibinders were amplified by polymerase chain reaction (PCR) using forward and reverse primers incorporating the promoter and RBS sequences required for LET production. PCR amplification was performed at an annealing temperature of 56 °C, and the amplified DNA products were used directly as LETs for subsequent CFPS without further purification.

Candidate minibinders were expressed using an *E. coli*-based CFPS system. Reactions were assembled in a final volume of 20 μL containing 1× salt solution (from a 15× stock), 1× master mix (from a 15× stock), 2 mM each amino acid, 33 mM phosphoenolpyruvate (PEP), 0.33 mM nicotinamide adenine dinucleotide (NAD), 0.27 mM coenzyme A (CoA), 4 mM oxalic acid, 1 mM putrescine, 1.5 mM spermidine, 57 mM HEPES, 0.1 mg/mL T7 RNA polymerase, and 40% (v/v) cell extract. Unpurified PCR-amplified linear DNA templates (1 μL) were added directly to the CFPS reactions, which were incubated overnight at 30 °C to allow protein expression. Following CFPS, the entire 20 μL reaction was diluted with 180 μL of 1× phosphate-buffered saline (PBS) and centrifuged at 3,000 rpm for 10 min to remove insoluble lysate. The clarified supernatant was used to evaluate minibinder binding and expression by ELISA. For each assay, 30 μL of clarified CFPS sample was added to individual wells in triplicate and supplemented with 70 μL of 1× PBS to obtain a final volume of 100 μL per well.

For the binding ELISA, high-binding 96-well plates (Corning) were coated with 1 μg of TEV-treated recombinant β-catenin in 100 μL PBS per well and incubated overnight at 4 °C. The plates were washed three times with PBS containing 0.05% Tween-20 (PBST) and blocked with 200 μL of 1% (w/v) bovine serum albumin (BSA) in PBS for 1 h at room temperature. Following blocking, the plates were washed three times with PBST, and 100 μL of diluted CFPS sample was added to each well. Samples were incubated for 2 h at room temperature with orbital shaking. The plates were subsequently washed three times with PBST and incubated with 100 μL of anti-His-HRP antibody (1:20,000 diluted, Abcam, ab1187) for 1 h at room temperature. Following five washes with PBS-T, 100 μL of 3,3′,5,5′-tetramethylbenzidine (TMB) substrate (1-Step Ultra TMB-ELISA; ThermoScientific) was added to each well and incubated for 10 min in the dark. The reaction was stopped by adding 100 μL of 2 M sulfuric acid, and absorbance was measured at 450 nm using a spectrophotometer. Minibinder expression was evaluated separately using a direct ELISA. Briefly, 100 μL of diluted CFPS sample was added directly to high-binding 96-well ELISA plates and incubated overnight at 4 °C to allow protein adsorption. The following day, the plates were washed with PBST, incubated with anti-His-HRP antibody (1:20,000 diluted, Abcam, ab1187) and Ultra-TMB as above to detect the expressed minibinders.

### Expression and purification of minibinders and β**-catenin**

Expression and purification of minibinders for subsequent assessment of purity, yield, and was completed as follows. First, a colony of BL21(DE3) cells carrying a pET28a plasmid encoding the relevant minibinder was inoculated in 25 mL of LB media supplemented with 50 μg/mL kanamycin (Kan) in a 125-mL culture flask and grown for 16 h at 37 °C with shaking at 250 rpm. Then, 20 mL of saturated starter culture was used to inoculate 1 L of LB in a 2.8–L culture flask supplemented with 50 μg/mL Kan. This culture was subsequently incubated at 37 °C with shaking at 250 rpm until the optical density at 600 nm (OD_600_) of the culture reached ~0.6–0.8, at which point protein expression was induced with 1 mM isopropyl-β-D-thiogalactoside (IPTG). Protein expression was allowed to proceed for 16–20 h at 30 °C with shaking at 250 rpm, after which cells were harvested via centrifugation by spinning at 8,000 x *g* and 4 °C for 15 min. Pellets were resuspended with 3 mL of Buffer W (100 mM Tris/HCl, 150 mM NaCl, 1 mM EDTA, pH to 8.0, sterile filtered) per 1 g of pellet and lysed on ice using a sonicator (QSonica) for 9 min with a duty cycle of 10s on : 10s off at an amplification of 40%. The soluble fraction was obtained as the supernatant after pelleting the lysate by spinning at 18,000 x *g* and 4 °C for 25 min. Meanwhile, 300 μL of Strep-Tactin 4Flow resin (IBA Life Sciences) was added to a 10-mL Poly-prep Chromatography Column (Bio-Rad) and equilibrated twice with 5 column volumes (CVs) of Buffer W. The supernatant was then sterile filtered and added to the column and allowed to flow through. Next, the column was washed five times with 1 to 2 CVs of Buffer W. To elute the protein, 0.8 CVs of Buffer BXT (100 mM Tris/HCl, 150 mM NaCl, 1 mM EDTA, 50 mM biotin, pH to 8.0, sterile filtered) was added to the column but not collected. Then the purified minibinders were collected by addition of 1.6 CVs of Buffer BXT followed by an additional 0.8 CVs of Buffer BXT. This collected elution material was subsequently concentrated to 100 uL using a 3k MWCO 0.5 mL Pierce Protein Concentrator (ThermoFisher) and dialyzed into 1× PBS overnight at 4 °C using 3.5k MWCO Slide-A-Lyzer™ MINI Dialysis Devices (ThermoFisher). The purified protein was subsequently stored at 4 °C until further use. If the column was reused, it was regenerated using 8 CVs of Buffer XT-R (3M MgCl_2_, sterile filtered), washed with 12 CVs of Buffer W, and stored at 4 °C in Buffer W. Purified proteins were quantified via BCA assay (ThermoFisher).

Expression and purification of β-catenin for binding assays was completed similarly to purification of minibinders, with the following changes: The 1–L subculture was induced with 0.5 mM IPTG at an OD_600_ of ~0.4–0.6 then incubated at 16 °C overnight. Pellets were resuspended in 2 mL per g pellet of resuspension buffer (50 mM NaH_2_PO_4_ 300 mM NaCl, pH 8.0, sterile filtered). After obtaining the sterile filtered soluble fraction, it was diluted with elution buffer (50 mM NaH_2_PO_4_ 300 mM NaCl, 500 mM imidazole, pH 8.0, sterile filtered) to a final imidazole concentration of 20 mM then purified and fractionated on a GE Healthcare fast protein liquid chromatography (FPLC) instrument using a HisTrap HP column (Cytivia) according to the manufacturer’s protocols. Fractions containing significant protein according to the UV-Vis trace were concentrated in a 30K MWCO 6–20 mL Pierce Protein Concentrator (ThermoFisher). Next, a PD-10 desalting column (Cytivia) was equilibrated with storage buffer (1× PBS, 1 mM DTT, 20% (v/v) glycerol) according to the manufacturer’s gravity column protocol. The concentrated β-catenin was then added to the column followed by a stacker of 0.75 mL of storage buffer. The β-catenin was eluted with 3.5 mL of storage buffer, aliquoted, flash frozen in liquid nitrogen, and stored at –20 °C. The concentration of β-catenin was subsequently determined by BCA assay (ThermoFisher) after thawing an aliquot.

### SDS-PAGE protein gel analysis

Protein samples that were purified via StrepXT gravity column purification were subsequently used for sodium dodecyl sulfate polyacrylamide gel electrophoresis (SDS-PAGE) analysis. These samples were solubilized in 10% (v/v) β-mercaptoethanol (BME) in 4× lithium dodecyl sulfate (LDS) buffer and subsequently heated at 98 °C for 10 min. They were then run on Bolt Bis-Tris Plus 12% 10-well protein gels (ThermoFisher) at 100V for 1.5 hr. Total protein staining of SDS-PAGE gels was accomplished by washing the gels three times in water for at least 5 min, staining for 1 h with Coomassie Blue G-250 stain (Bio-Rad), then destaining for at least 1 h in water. Gels were imaged using the Chemidoc MP (Bio-Rad) and purity was determined using band densitometry calculated in ImageLab 6.1 (Bio-Rad).

### EC_**50**_ **ELISA**

The EC_50_ affinity measurement ELISA was conducted as follows: 96-well plates (Corning) were coated with purified β-catenin (2 μg/ml) in 1× PBS (pH 7.4) at a volume of 50 μl per well at 4 °C overnight. Next, the plates were washed four times with 300 uL of 1× PBS with 0.1% (v/v) Tween 20 (PBST), hand decanting each time, and then blocked with 300 μL of blocking buffer (5% BSA in 1× TBST) per well overnight at 4 °C. Purified minibinders in 1× PBS were biotinylated using EZ-Link™ Sulfo NHS-LC-LC-Biotin (Thermo Fisher Scientific) following the manufacturer’s protocol using a MR of 10 and resuspending the reagent in DMSO. The resulting biotinylated products were then quenched with a final concentration of 100 mM Tris and diluted into 1× PBS before being serially diluted using 1× PBS and added to the ELISA plates for 2 h at RT with gentle shaking. The plates were then washed four times with 1× PBST and incubated for 1 h shaking at RT with 100 μL of Extravidin-Peroxidase (diluted 1:20,000; Thermo Fisher Scientific, E2886). After an additional four washes with 1× PBST, 100 μL of 3,3′-5,5′-tetramethylbenzidine substrate (1-Step Ultra TMB-ELISA; Thermo Fisher Scientific) was added to each well, and the plates were incubated at room temperature in darkness. The reaction was quenched by adding 100 μL of 2 M H_2_SO_4_, and absorbance was quantified at a wavelength of 450 nm using a FilterMax F5 microplate spectrophotometer (Agilent).

### BLI

Binding affinity of the purified minibinders was measured by BLI using an Octet RH16 instrument (Sartorius) at 30 °C shaking at 1000 rpm. Kinetic analysis was performed using streptavidin biosensor tips (Sartorius, 18-5020) as follows: (i) initial sensor equilibration: 150-s immersion in Octet buffer (1× PBS containing 0.2-μm filtered 0.1% BSA and 0.02% (v/v) Tween 20); (ii) loading: 150-s immersion in Octet buffer supplemented with biotinylated minibinders (3 μg/ml) except for one well which was used with the reference sample; (iii) post-loading equilibration: 150-s immersion in Octet buffer; (iv) association: 1000-s immersion in solution with β-catenin at concentrations of 4000, 2000, 1000, 500, 250, and 125 nM, plus 1000 nM for the reference sample and 0 nM for the reference sensor; (v) dissociation: 1000-s immersion in Octet buffer using the wells previously used for post-loading equilibration. All kinetic data were analyzed using Octet Analysis Studio (Sartorius).

### Cell culture and transfection

HEK293T cells were maintained in Dulbecco’s modified Eagle medium (DMEM; Gibco) supplemented with 10% serum (HyClone) at 37 °C in a humidified atmosphere containing 5% Co□. Cells were seeded at a density of 2.5 × 10□ cells/mL 24 h before transfection. The next day, cells were transfected with plasmid DNA (pDNA) or CTNNB1 siRNA (Cell Signaling Technology) using jetPRIME transfection reagent (Polyplus-transfection; VWR) according to the manufacturer’s instructions. Briefly, pDNA was diluted in jetPRIME buffer, and jetPRIME reagent was added at a pDNA:reagent ratio of 1:2. For siRNA transfection, siRNA was diluted in jetPRIME buffer followed by the addition of jetPRIME reagent to achieve a final concentration of 50 nM per well. In both cases, the mixtures were briefly vortexed and incubated for 10 min at room temperature before being added dropwise to the cells. Cells were incubated for 48 h after transfection before subsequent analysis.

### Immunoblot analysis

HEK293T cells transfected with pcDNA3-based plasmids encoding CHIPΔTPR ^24^, B3-CHIPΔTPR, or C4-CHIPΔTPR were harvested at 48 h post-transfection and washed with PBS. Cytosolic cellular fractions were isolated following the cultured cell fractionation protocol ^49^. Specifically, ice-cold lysis buffer A containing 150 mM NaCl, 50 mM HEPES (pH 7.4), 25 μg/mL digitonin (Sigma), 1 M hexylene glycol (Sigma), and 1% (v/v) protease inhibitor (Thermo Fisher Scientific) was added to the pellets, and samples were incubated for 10 min at 4 °C on an end-over-end rotator, followed by centrifugation at 2,000 × *g* for 10 min at 4 °C. Protein concentrations were determined using the Pierce BCA Protein Assay Kit (Thermo Fisher Scientific), and equal amounts of total protein were resolved by SDS-PAGE using Bolt Bis-Tris 4–12% protein gels (Thermo Fisher Scientific). Proteins were transferred to polyvinylidene fluoride (PVDF) membranes (Millipore) and blocked with 5% (w/v) nonfat dry milk (Carnation) in 1× Tris-buffered saline with 0.05% (v/v) Tween 20 (TBST) for 1 h at room temperature. Membranes were incubated with anti-β-catenin (Cell Signaling Technology, cat # 37447S; 1:2000) and anti-GAPDH (Abcam, cat # ab59164; 1:5000) antibodies overnight at 4 °C. Following three 10-minute washes with TBST, membranes were incubated with goat anti-mouse IgG Fc (HRP) secondary antibody (Abcam, cat # ab97265; 1:5000) for 1 h at room temperature. Protein bands were detected using Clarity Western ECL Substrate (Bio-Rad) and imaged using a ChemiDoc MP imager (Bio-Rad). Band intensities were quantified using ImageJ software (version 1.37v) as described at https://imagej.net. β-catenin band intensities were normalized to the corresponding GAPDH band intensities and expressed relative to the CHIPΔTPR control.

### TOPFlash assay

HEK293T cells were seeded in 12-well plates at a density of 2.5 × 10LJ cells/mL 24 h before transfection. The next day, cells were co-transfected with the M50 Super 8x TOPFlash plasmid (Addgene, plasmid #12456) or the M51 Super 8x FOPFlash plasmid (TOPFlash mutant; Addgene, plasmid #12457), the pCMV-Renilla plasmid, and a pcDNA3 plasmid encoding CHIPΔTPR, B3-CHIPΔTPR, or C4-CHIPΔTPR. Each well received 1.5 μg of total pDNA at a TOPFlash/FOPFlash:Renilla:pcDNA3 mass ratio of 1:0.1:3 using jetPRIME transfection reagent. At 48 h after transfection, cells were detached and lysed in the passive lysis buffer supplied with the dual-luciferase reporter kit (Promega). Aliquots of each lysate (20 μL per well) were added in technical triplicate to a white 96-well plate. Firefly and Renilla luminescence signals were measured sequentially using the dual-luciferase reporter kit (Promega) according to the manufacturer’s instructions on a microplate reader (Tecan Spark). Firefly luciferase signals were normalized to the corresponding Renilla luciferase signals, and expression was measured relative to the CHIPΔTPR control. For TOPFlash analysis of siRNA-treated cells, the same protocol was followed, except that CTNNB1 siRNA was transfected at a final concentration of 50 nM in place of plasmid DNA, and signals were likewise expressed relative to the CHIPΔTPR control.

## Acknowledgements.

This work was supported by the Defense Advanced Research Projects Agency (DARPA contract W911NF-23-2-0039 to M.P.D.), the Defense Threat Reduction Agency (grant HDTRA1-20-10004 and HDTRA1-26-1004 to M.P.D.), the National Institutes of Health (grant R35GM155380 to W.K.M.L.), and the National Science Foundation (grants CBET-1605242 and CBET-1936823 to M.P.D.). E.J.B. was supported by an NIH/NIGMS Chemical Biology Interface Training Grant (T32GM138826) and an NSF Graduate Research Fellowship (DGE-2139899).

## REFERENCES

(1) Tsai, J. M.; Nowak, R. P.; Ebert, B. L.; Fischer, E. S. Targeted Protein Degradation: From Mechanisms to Clinic. Nat. Rev. Mol. Cell Biol. 2024, 25 (9), 740–757. 10.1038/s41580-024-00729-9

(2) Dang, C. V.; Reddy, E. P.; Shokat, K. M.; Soucek, L. Drugging the “undruggable” Cancer Targets. Nat. Rev. Cancer 2017, 17 (8), 502–508. 10.1038/nrc.2017.36

(3) Henley, M. J.; Koehler, A. N. Advances in Targeting ‘Undruggable’ Transcription Factors with Small Molecules. Nat. Rev. Drug Discov. 2021, 20 (9), 669–688. 10.1038/s41573-021-00199-0

(4) Lai, A. C.; Crews, C. M. Induced Protein Degradation: An Emerging Drug Discovery Paradigm. Nat. Rev. Drug Discov. 2017, 16 (2), 101–114. 10.1038/nrd.2016.211

(5) Ye, T.; Alamgir, A.; Robertus, C. M.; Colina, D.; Monticello, C.; Donahue, T. C.; Bandi, C.; Hong, L.; Vincoff, S.; Goel, S.; Lam, K.; Heyes, J.; Putnam, D.; Alabi, C. A.; Chatterjee, P.; DeLisa, M. P. Programmable Protein Degraders Enable Selective Knockdown of Pathogenic β-Catenin Subpopulations in Vitro and in Vivo. Sci. Adv. 2026, 12 (35), eaec8271. 10.1126/sciadv.aec8271

(6) Lim, S.; Khoo, R.; Peh, K. M.; Teo, J.; Chang, S. C.; Ng, S.; Beilhartz, G. L.; Melnyk, R. A.; Johannes, C. W.; Brown, C. J.; Lane, D. P.; Henry, B.; Partridge, A. W. bioPROTACs as Versatile Modulators of Intracellular Therapeutic Targets Including Proliferating Cell Nuclear Antigen (PCNA). Proc. Natl. Acad. Sci. 2020, 117 (11), 5791–5800. 10.1073/pnas.1920251117

(7) Portnoff, A. D.; Stephens, E. A.; Varner, J. D.; DeLisa, M. P. Ubiquibodies, Synthetic E3 Ubiquitin Ligases Endowed with Unnatural Substrate Specificity for Targeted Protein Silencing. J. Biol. Chem. 2014, 289 (11), 7844–7855. 10.1074/jbc.M113.544825

(8) Yang, W.; Wang, S.; Lee, G. R.; Zhang, J. Z.; Courbet, A.; Juergens, D.; Wang, X.; Schlichthaerle, T.; Abedi, M.; Ragotte, R.; An, L.; Kalvet, I.; Pellock, S.; Mihaljevic, L.; Glasscock, C.; Pillai, A.; Broerman, A.; Ennist, N.; Haefner, E.; McNamara-Bordewick, N.; Haydon, I.; Stewart, L.; Bhardwaj, G.; Baker, D. The Past, Present and Future of de Novo Protein Design. Nature 2026, 652 (8112), 1139–1152. 10.1038/s41586-026-10328-7

(9) Cao, L.; Coventry, B.; Goreshnik, I.; Huang, B.; Sheffler, W.; Park, J. S.; Jude, K. M.; Marković, I.; Kadam, R. U.; Verschueren, K. H. G.; Verstraete, K.; Walsh, S. T. R.; Bennett, N.; Phal, A.; Yang, A.; Kozodoy, L.; DeWitt, M.; Picton, L.; Miller, L.; Strauch, E.-M.; DeBouver, N. D.; Pires, A.; Bera, A. K.; Halabiya, S.; Hammerson, B.; Yang, W.; Bernard, S.; Stewart, L.; Wilson, I. A.; Ruohola-Baker, H.; Schlessinger, J.; Lee, S.; Savvides, S. N.; Garcia, K. C.; Baker, D. Design of Protein-Binding Proteins from the Target Structure Alone. Nature 2022, 605 (7910), 551–560. 10.1038/s41586-022-04654-9

(10) Chevalier, A.; Silva, D.-A.; Rocklin, G. J.; Hicks, D. R.; Vergara, R.; Murapa, P.; Bernard, S. M.; Zhang, L.; Lam, K.-H.; Yao, G.; Bahl, C. D.; Miyashita, S.-I.; Goreshnik, I.; Fuller, J. T.; Koday, M. T.; Jenkins, C. M.; Colvin, T.; Carter, L.; Bohn, A.; Bryan, C. M.; Fernández-Velasco, D. A.; Stewart, L.; Dong, M.; Huang, X.; Jin, R.; Wilson, I. A.; Fuller, D. H.; Baker, D. Massively Parallel de Novo Protein Design for Targeted Therapeutics. Nature 2017, 550 (7674), 74–79. 10.1038/nature23912

(11) Vázquez Torres, S.; Benard Valle, M.; Mackessy, S. P.; Menzies, S. K.; Casewell, N. R.; Ahmadi, S.; Burlet, N. J.; Muratspahić, E.; Sappington, I.; Overath, M. D.; Rivera-de-Torre, E.; Ledergerber, J.; Laustsen, A. H.; Boddum, K.; Bera, A. K.; Kang, A.; Brackenbrough, E.; Cardoso, I. A.; Crittenden, E. P.; Edge, R. J.; Decarreau, J.; Ragotte, R. J.; Pillai, A. S.; Abedi, M.; Han, H. L.; Gerben, S. R.; Murray, A.; Skotheim, R.; Stuart, L.; Stewart, L.; Fryer, T. J. A.; Jenkins, T. P.; Baker, D. De Novo Designed Proteins Neutralize Lethal Snake Venom Toxins. Nature 2025, 639 (8053), 225–231. 10.1038/s41586-024-08393-x

(12) Muratspahić, E.; Feldman, D.; Kim, D. E.; Qu, X.; Bratovianu, A.-M.; Rivera-Sánchez, P.; Voss, J. H.; Hertz, E. P. T.; Jeppesen, M.; Dimitri, F.; Sakamoto, K.; Nallathambi, A.; Peceli, P.; Cao, J.; Cary, B. P.; Belousoff, M. J.; Keov, P.; Trinh, P. N. H.; Chen, Q.; Ren, Y.; Fine, J.; Mishra, S.; Dalal, A.; Sinha, S.; Banerjee, R.; Ganguly, M.; Karuppusamy, K. V.; Sappington, I.; Schlichthaerle, T.; Zhang, J. Z.; Pillai, A.; Coventry, B.; Mihaljević, L.; Bauer, M. S.; Torres, S. V.; Motmaen, A.; Lee, G. R.; Tran, L.; Wang, X.; Goreshnik, I.; Vafeados, D. K.; Svendsen, J. E.; Hosseinzadeh, P.; Lindegaard, N.; Brandt, M.; Waltenspühl, Y.; Deibler, K.; Deweid, L.; Bennett, A.; Schöppe, J.; Dong, T.; Yan, X.; Oostdyk, L.; Cao, W.; Anantharaman, L.; Weisser, J. J.; Bastlund, J. F.; Bundgaard, C.; Asuni, A. A.; English, J. G.; Stewart, L. J.; Halloran, L.; Spangler, J. B.; Lieber, A.; Shukla, A. K.; Sexton, P. M.; Roth, B. L.; Krumm, B. E.; Wootten, D.; Tate, C. G.; Norn, C.; Baker, D. De Novo Design of Miniproteins Targeting GPCRs. Nature 2026, 656 (8129), 1044–1053. 10.1038/s41586-026-10656-8

(13) Cao, L.; Goreshnik, I.; Coventry, B.; Case, J. B.; Miller, L.; Kozodoy, L.; Chen, R. E.; Carter, L.; Walls, A. C.; Park, Y.-J.; Strauch, E.-M.; Stewart, L.; Diamond, M. S.; Veesler, D.; Baker, D. De Novo Design of Picomolar SARS-CoV-2 Miniprotein Inhibitors. Science 2020, 370 (6515), 426–431. 10.1126/science.abd9909

(14) Park, J. S.; Choi, J.; Cao, L.; Mohanty, J.; Suzuki, Y.; Park, A.; Baker, D.; Schlessinger, J.; Lee, S. Isoform-Specific Inhibition of FGFR Signaling Achieved by a de-Novo-Designed Mini-Protein. Cell Rep. 2022, 41 (4), 111545. 10.1016/j.celrep.2022.111545

(15) Lv, X.; Zhang, Y.; Sun, K.; Yang, Q.; Luo, J.; Tao, L.; Lu, P. De Novo Design of Mini-Protein Binders Broadly Neutralizing Clostridioides Difficile Toxin B Variants. Nat. Commun. 2024, 15 (1), 8521. 10.1038/s41467-024-52582-1

(16) Gokce-Alpkilic, G.; Huang, B.; Liu, A.; Kreuk, L. S. M.; Wang, Y.; Adebomi, V.; Bueso, Y. F.; Bera, A. K.; Kang, A.; Gerben, S. R.; Rettie, S.; Vafeados, D. K.; Roullier, N.; Goreshnik, I.; Li, X.; Baker, D.; Woodward, J. J.; Mougous, J. D.; Bhardwaj, G. De Novo Design of High-Affinity Miniprotein Binders Targeting Francisella Tularensis Virulence Factor. Angew. Chem. Int. Ed. 2025, 64 (52), e202516058. 10.1002/anie.202516058

(17) Yang, W.; Hicks, D. R.; Ghosh, A.; Schwartze, T. A.; Conventry, B.; Goreshnik, I.; Allen, A.; Halabiya, S. F.; Kim, C. J.; Hinck, C. S.; Lee, D. S.; Bera, A. K.; Li, Z.; Wang, Y.; Schlichthaerle, T.; Cao, L.; Huang, B.; Garrett, S.; Gerben, S. R.; Rettie, S.; Heine, P.; Murray, A.; Edman, N.; Carter, L.; Stewart, L.; Almo, S. C.; Hinck, A. P.; Baker, D. Design of High-Affinity Binders to Immune Modulating Receptors for Cancer Immunotherapy. Nat. Commun. 2025, 16 (1), 2001. 10.1038/s41467-025-57192-z

(18) Huang, B.; Abedi, M.; Ahn, G.; Coventry, B.; Sappington, I.; Tang, C.; Wang, R.; Schlichthaerle, T.; Zhang, J. Z.; Wang, Y.; Goreshnik, I.; Chiu, C. W.; Chazin-Gray, A.; Chan, S.; Gerben, S.; Murray, A.; Wang, S.; O’Neill, J.; Yi, L.; Yeh, R.; Misquith, A.; Wolf, A.; Tomasovic, L. M.; Piraner, D. I.; Duran Gonzalez, M. J.; Bennett, N. R.; Venkatesh, P.; Ahlrichs, M.; Dobbins, C.; Yang, W.; Wang, X.; Sahtoe, D. D.; Vafeados, D.; Mout, R.; Shivaei, S.; Cao, L.; Carter, L.; Stewart, L.; Spangler, J. B.; Roybal, K. T.; Greisen, P. J.; Li, X.; Bernardes, G. J. L.; Bertozzi, C. R.; Baker, D. Designed Endocytosis-Inducing Proteins Degrade Targets and Amplify Signals. Nature 2025, 638 (8051), 796–804. 10.1038/s41586-024-07948-2

(19) Weinberg, Z. Y.; Soliman, S. S.; Kim, M. S.; Shah, D. H.; Chen, I. P.; Ott, M.; Lim, W. A.; El-Samad, H. De Novo-Designed Minibinders Expand the Synthetic Biology Sensing Repertoire. eLife 2024, 13. 10.7554/eLife.96154.2

(20) Kortemme, T. De Novo Protein Design—From New Structures to Programmable Functions. Cell 2024, 187 (3), 526–544. 10.1016/j.cell.2023.12.028

(21) Chatterjee, P.; Ponnapati, M.; Kramme, C.; Plesa, A. M.; Church, G. M.; Jacobson, J. M. Targeted Intracellular Degradation of SARS-CoV-2 via Computationally Optimized Peptide Fusions. Commun. Biol. 2020, 3 (1), 715. 10.1038/s42003-020-01470-7

(22) Brixi, G.; Ye, T.; Hong, L.; Wang, T.; Monticello, C.; Lopez-Barbosa, N.; Vincoff, S.; Yudistyra, V.; Zhao, L.; Haarer, E.; Chen, T.; Pertsemlidis, S.; Palepu, K.; Bhat, S.; Christopher, J.; Li, X.; Liu, T.; Zhang, S.; Petersen, L.; DeLisa, M. P.; Chatterjee, P. SaLT&PepPr Is an Interface-Predicting Language Model for Designing Peptide-Guided Protein Degraders. Commun. Biol. 2023, 6 (1), 1081. 10.1038/s42003-023-05464-z

(23) Bhat, S.; Palepu, K.; Hong, L.; Mao, J.; Ye, T.; Iyer, R.; Zhao, L.; Chen, T.; Vincoff, S.; Watson, R.; Wang, T. Z.; Srijay, D.; Kavirayuni, V. S.; Kholina, K.; Goel, S.; Vure, P.; Deshpande, A. J.; Soderling, S. H.; DeLisa, M. P.; Chatterjee, P. De Novo Design of Peptide Binders to Conformationally Diverse Targets with Contrastive Language Modeling. Sci. Adv. 2025, 11 (4), eadr8638. 10.1126/sciadv.adr8638

(24) Ye, T.; Alamgir, A.; Robertus, C. M.; Colina, D.; Monticello, C.; Donahue, T. C.; Bandi, C.; Hong, L.; Vincoff, S.; Goel, S.; Lam, K.; Heyes, J.; Putnam, D.; Alabi, C. A.; Chatterjee, P.; DeLisa, M. P. Programmable Protein Degraders Enable Selective Knockdown of Pathogenic β-Catenin Subpopulations in Vitro and in Vivo. Sci. Adv. 2026, 12 (35), eaec8271. 10.1126/sciadv.aec8271

(25) Chen, L. T.; Quinn, Z.; Dumas, M.; Peng, C.; Hong, L.; Lopez-Gonzalez, M.; Mestre, A.; Watson, R.; Vincoff, S.; Zhao, L.; Wu, J.; Stavrand, A.; Schaepers-Cheu, M.; Wang, T. Z.; Srijay, D.; Monticello, C.; Vure, P.; Pulugurta, R.; Pertsemlidis, S.; Kholina, K.; Goel, S.; DeLisa, M. P.; Chi, J.-T. A.; Truant, R.; Aguilar, H. C.; Chatterjee, P. Target Sequence-Conditioned Design of Peptide Binders Using Masked Language Modeling. Nat. Biotechnol. 2026, 44 (6), 1002–1010. 10.1038/s41587-025-02761-2

(26) Hunt, A. C.; Vögeli, B.; Hassan, A. O.; Guerrero, L.; Kightlinger, W.; Yoesep, D. J.; Krüger, A.; DeWinter, M.; Diamond, M. S.; Karim, A. S.; Jewett, M. C. A Rapid Cell-Free Expression and Screening Platform for Antibody Discovery. Nat. Commun. 2023, 14 (1), 3897. 10.1038/s41467-023-38965-w

(27) Landwehr, G. M.; Bogart, J. W.; Magalhaes, C.; Hammarlund, E. G.; Karim, A. S.; Jewett, M. C. Accelerated Enzyme Engineering by Machine-Learning Guided Cell-Free Expression. Nat. Commun. 2025, 16 (1), 865. 10.1038/s41467-024-55399-0

(28) Silverman, A. D.; Karim, A. S.; Jewett, M. C. Cell-Free Gene Expression: An Expanded Repertoire of Applications. Nat. Rev. Genet. 2020, 21 (3), 151–170. 10.1038/s41576-019-0186-3

(29) Lange, B.; Godin, R.; Schnicker, N. J.; Wadas, T. J.; Reuel, N. F. Rapid Affinity Characterization of Cell-Free Expressed Nanobodies Directly in Lysate with Biolayer Interferometry via Antigen-Immobilized Biosensors. ACS Synth. Biol. 2026. 10.1021/acssynbio.6c00062

(30) Zhang, H.; Chen, M.; Li, Y.; He, Q.; Tu, Z. A Rapid Cell-Free Expression Platform for Screening of Nanobody Mutants. Biotechnol. Bioprocess Eng. 2026. 10.1007/s12257-026-00273-7

(31) Clark-ElSayed, A.; Harrison, I. M.; Olsen, M. L.; Lazar, J. T.; Jewett, M. C.; Ellington, A. D. LDBT Instead of DBTL: Combining Machine Learning and Rapid Cell-Free Testing. Nat. Commun. 2025, 16 (1), 9782. 10.1038/s41467-025-65281-2

(32) Capin, J.; Mayonove, P.; DeVisch, A.; Becher, A.; Ngo, G.; Courbet, A.; Ragotte, R. J.; Cohen Gonsaud, M.; Espeut, J.; Bonnet, J. CF2H: A Cell-Free Two-Hybrid Platform for Rapid Protein Binder Screening. Nat. Commun. 2026, 17 (1), 3724. 10.1038/s41467-026-69741-1

(33) Cui, C.; Zhou, X.; Zhang, W.; Qu, Y.; Ke, X. Is β-Catenin a Druggable Target for Cancer Therapy? Trends Biochem. Sci. 2018, 43 (8), 623–634. 10.1016/j.tibs.2018.06.003

(34) Watson, J. L.; Juergens, D.; Bennett, N. R.; Trippe, B. L.; Yim, J.; Eisenach, H. E.; Ahern, W.; Borst, A. J.; Ragotte, R. J.; Milles, L. F.; Wicky, B. I. M.; Hanikel, N.; Pellock, S. J.; Courbet, A.; Sheffler, W.; Wang, J.; Venkatesh, P.; Sappington, I.; Torres, S. V.; Lauko, A.; De Bortoli, V.; Mathieu, E.; Ovchinnikov, S.; Barzilay, R.; Jaakkola, T. S.; DiMaio, F.; Baek, M.; Baker, D. De Novo Design of Protein Structure and Function with RFdiffusion. Nature 2023, 620 (7976), 1089–1100. 10.1038/s41586-023-06415-8

(35) Nusse, R.; Clevers, H. Wnt/β-Catenin Signaling, Disease, and Emerging Therapeutic Modalities. Cell 2017, 169 (6), 985–999. 10.1016/j.cell.2017.05.016

(36) Schinn, S.-M.; Broadbent, A.; Bradley, W. T.; Bundy, B. C. Protein Synthesis Directly from PCR: Progress and Applications of Cell-Free Protein Synthesis with Linear DNA. New Biotechnol. 2016, 33 (4), 480–487. 10.1016/j.nbt.2016.04.002

(37) Portnoff, A. D.; Stephens, E. A.; Varner, J. D.; DeLisa, M. P. Ubiquibodies, Synthetic E3 Ubiquitin Ligases Endowed with Unnatural Substrate Specificity for Targeted Protein Silencing. J. Biol. Chem. 2014, 289 (11), 7844–7855. 10.1074/jbc.M113.544825

(38) Cadigan, K. M.; Waterman, M. L. TCF/LEFs and Wnt Signaling in the Nucleus. Cold Spring Harb. Perspect. Biol. 2012, 4 (11), a007906. 10.1101/cshperspect.a007906

(39) Cong, F.; Zhang, J.; Pao, W.; Zhou, P.; Varmus, H. A Protein Knockdown Strategy to Study the Function of Beta-Catenin in Tumorigenesis. BMC Mol. Biol. 2003, 4, 10. 10.1186/1471-2199-4-10

(40) Veeman, M. T.; Slusarski, D. C.; Kaykas, A.; Louie, S. H.; Moon, R. T. Zebrafish Prickle, a Modulator of Noncanonical Wnt/Fz Signaling, Regulates Gastrulation Movements. Curr. Biol. CB 2003, 13 (8), 680–685. 10.1016/s0960-9822(03)00240-9

(41) Bennett, N. R.; Coventry, B.; Goreshnik, I.; Huang, B.; Allen, A.; Vafeados, D.; Peng, Y. P.; Dauparas, J.; Baek, M.; Stewart, L.; DiMaio, F.; De Munck, S.; Savvides, S. N.; Baker, D. Improving de Novo Protein Binder Design with Deep Learning. Nat. Commun. 2023, 14 (1), 2625. 10.1038/s41467-023-38328-5

(42) Stankiewicz, M.; Nikolay, R.; Rybin, V.; Mayer, M. P. CHIP Participates in Protein Triage Decisions by Preferentially Ubiquitinating Hsp70-Bound Substrates. FEBS J. 2010, 277 (16), 3353–3367. 10.1111/j.1742-4658.2010.07737.x

(43) Bondeson, D. P.; Mares, A.; Smith, I. E. D.; Ko, E.; Campos, S.; Miah, A. H.; Mulholland, K. E.; Routly, N.; Buckley, D. L.; Gustafson, J. L.; Zinn, N.; Grandi, P.; Shimamura, S.; Bergamini, G.; Faelth-Savitski, M.; Bantscheff, M.; Cox, C.; Gordon, D. A.; Willard, R. R.; Flanagan, J. J.; Casillas, L. N.; Votta, B. J.; den Besten, W.; Famm, K.; Kruidenier, L.; Carter, P. S.; Harling, J. D.; Churcher, I.; Crews, C. M. Catalytic in Vivo Protein Knockdown by Small-Molecule PROTACs. Nat. Chem. Biol. 2015, 11 (8), 611–617. 10.1038/nchembio.1858

(44) VanDyke, D.; Taylor, J. D.; Kaeo, K. J.; Hunt, J.; Spangler, J. B. Biologics-Based Degraders — an Expanding Toolkit for Targeted-Protein Degradation. Curr. Opin. Biotechnol. 2022, 78, 102807. 10.1016/j.copbio.2022.102807

(45) Qian, J.; Milles, L. F.; Wicky, B. I. M.; Ragotte, R. J.; Motmaen, A.; Borst, A. J.; Skotheim, R.; Ols, S.; Coventry, B.; Li, X.; Kibler, R. D.; Goreshnik, I.; Expòsit, M.; Loré, K.; Stewart, L.; Baker, D. Accelerating Protein Design by Scaling Experimental Characterization. Nat. Commun. 2026, 17 (1), 9945. 10.1038/s41467-026-76740-9

(46) Bienz, M. Beta-Catenin: A Pivot between Cell Adhesion and Wnt Signalling. Curr. Biol. CB 2005, 15 (2), R64–67. 10.1016/j.cub.2004.12.058

(47) Buechel, D.; Sugiyama, N.; Rubinstein, N.; Saxena, M.; Kalathur, R. K. R.; Lüönd, F.; Vafaizadeh, V.; Valenta, T.; Hausmann, G.; Cantù, C.; Basler, K.; Christofori, G. Parsing β-Catenin’s Cell Adhesion and Wnt Signaling Functions in Malignant Mammary Tumor Progression. Proc. Natl. Acad. Sci. U. S. A. 2021, 118 (34), e2020227118. 10.1073/pnas.2020227118

(48) Varadi, M.; Anyango, S.; Deshpande, M.; Nair, S.; Natassia, C.; Yordanova, G.; Yuan, D.; Stroe, O.; Wood, G.; Laydon, A.; Žídek, A.; Green, T.; Tunyasuvunakool, K.; Petersen, S.; Jumper, J.; Clancy, E.; Green, R.; Vora, A.; Lutfi, M.; Figurnov, M.; Cowie, A.; Hobbs, N.; Kohli, P.; Kleywegt, G.; Birney, E.; Hassabis, D.; Velankar, S. AlphaFold Protein Structure Database: Massively Expanding the Structural Coverage of Protein-Sequence Space with High-Accuracy Models. Nucleic Acids Res. 2022, 50 (D1), D439–D444. 10.1093/nar/gkab1061

(49) Baghirova, S.; Hughes, B. G.; Hendzel, M. J.; Schulz, R. Sequential Fractionation and Isolation of Subcellular Proteins from Tissue or Cultured Cells. MethodsX 2015, 2, 440–445. 10.1016/j.mex.2015.11.001

