## Supplementary Information for "Cell-free identification of *de novo* designed minibinders as building blocks for targeted protein degraders"

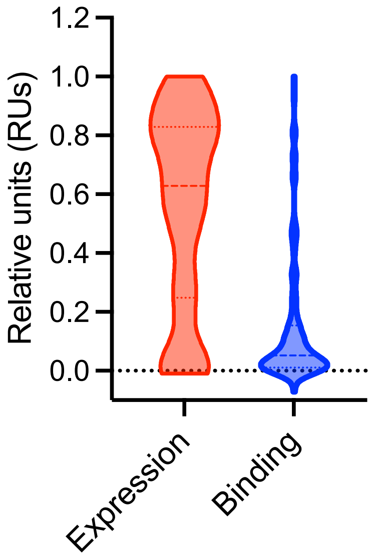


**Supplementary Figure 1. Expression and binding distribution of minibinders as determined by CFPS-ELISA.** Violin plot of relative minibinder expression levels and binding activity determined by ELISA analysis of diluted CFPS reactions. Expression levels were obtained by direct ELISA using diluted CFPS reactions for coating and subsequent minibinder detection using an anti-His-HRP antibody (Abcam). Binding activity was determined by ELISA using plates coated with recombinantly expressed β-catenin and probed with Extravidin-HRP (Sigma-Aldrich) to specifically detect miniprotein binding. Expression and binding values were obtained by subtracting the mean absorbance at 450 nm of blank wells from each well on the plate and then normalizing to a maximum value of 1 for expression and binding data separately. Relative units (RUs) for each well represent the mean of replicate CFPS reactions (*n* = 3).

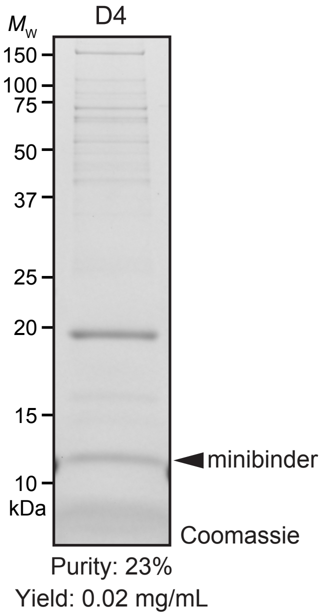


**Supplementary Figure 2. Purity and yield of purified minibinder D4.** SDS-PAGE analysis of minibinder D4 following expression in recombinant E. coli and purification by StrepXT affinity chromatography. Gel stained with Coomassie Brilliant Blue G250 (Bio-Rad). Protein purity was determined by densitometry analysis using ImageLab 6.1 (Bio-Rad) and protein yield was calculated by normalizing total protein content determined from BCA assay by the purity.

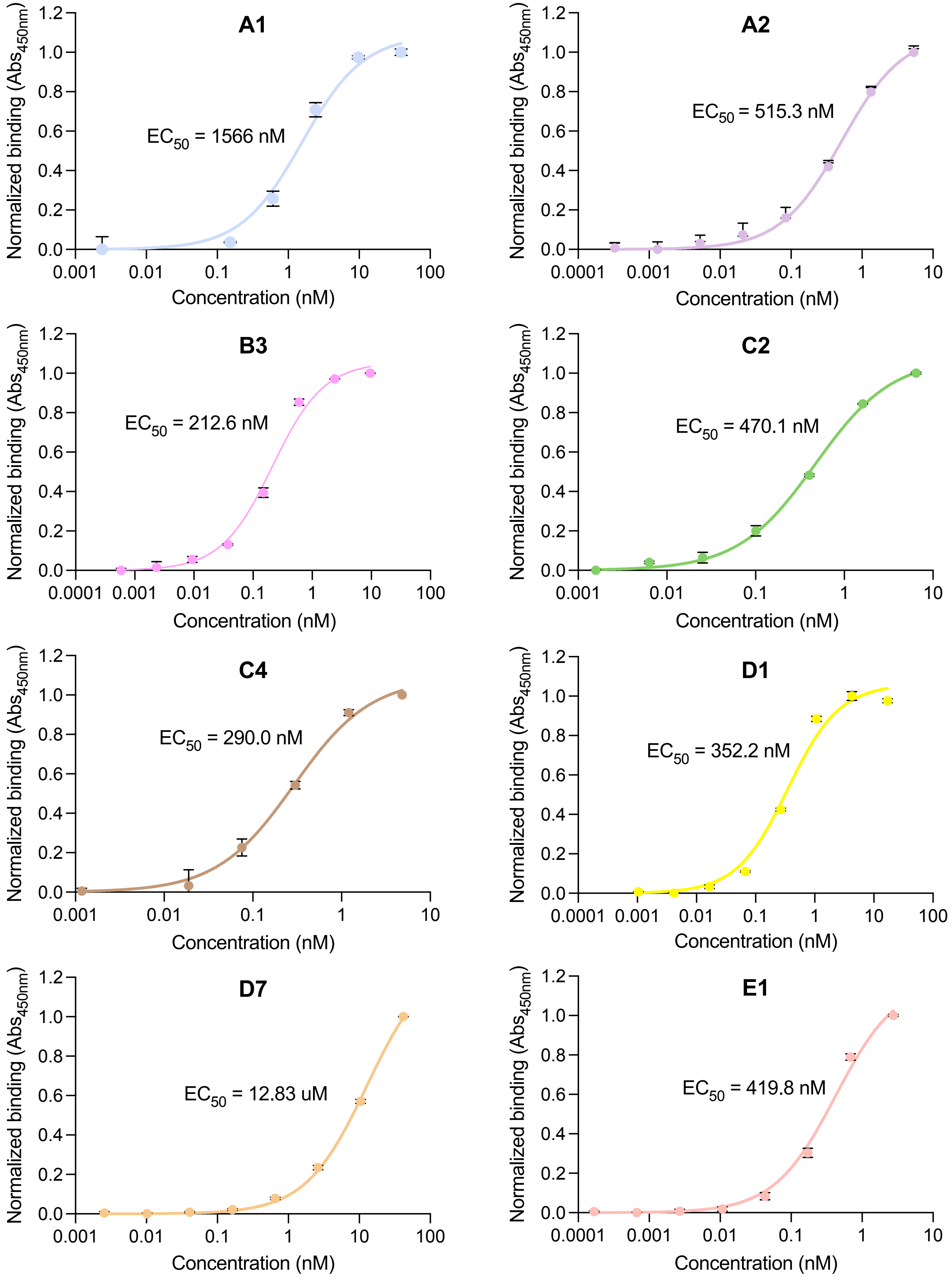


**Supplementary Figure 3. Binding analysis of downselected minibinders.** Individual binding curves for each minibinder following expression in recombinant *E. coli* and purification by StrepXT affinity chromatography. Binding activity was determined by ELISA with recombinant β-catenin as the immobilized antigen. Data are the average of technical replicates ($n$ = 2) ± SD. $EC_{50}$ values were obtained from curve fit analysis using GraphPad Prism for macOS (version 11.1.0).

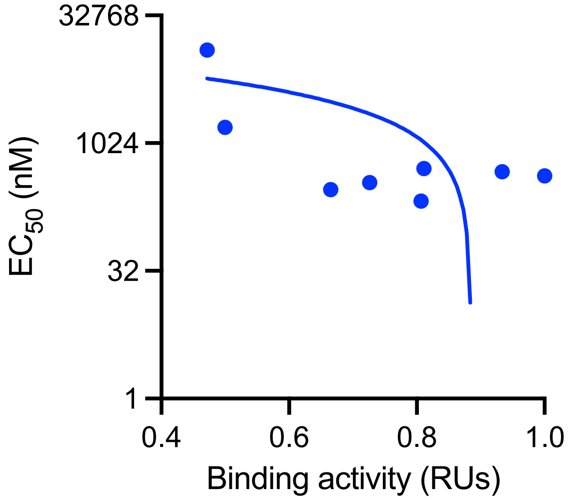


**Supplementary Figure 4. Relationship between apparent affinity ($EC_{50}$) and relative binding activity.** Relative miniprotein binding levels were obtained by ELISA analysis of diluted CFPS reactions using plates coated with recombinantly expressed β-catenin and probed with Extravidin-HRP to specifically detect miniprotein binding. Binding RUs are the average of biological replicates ($n$ = 3). $EC_{50}$ values for purified minibinders was determined by ELISA with recombinant β-catenin as the immobilized antigen and subsequent curve fit analysis using GraphPad Prism for macOS (version 11.1.0).
